# Clonal propagation of *Osyris lanceolata* through air layering at Bazawit Hill, Northern Ethiopia. An Endangered Medicinal and sandal wood Plant in East Africa

**DOI:** 10.1101/2020.03.23.002907

**Authors:** Adugnaw Admas, Smegnew Melese, Amare Genetu, Berhane Kidane, Zewdu Yilma, Melaku Admasu, Tesaka Misga

**Affiliations:** Ethiopian Enviroment and Forest Research Institute; Oganization for rehablitations and devlopment of Ahmara

**Keywords:** Osyris lanceolata, air layering, IBA application, rooting success

## Abstract

In the recent years medicinal and other economical important plants are getting attention due to the presence of therapeutically important active contents; however over exploitation and propagation problems are the major concern for conservation of several econmically important plant species. Among economical imprortant plants an attempt has been made to develop a propagation method for African sandalwood (*Osyris lanceolata*) by air layering approach aiming at providing an alternative propagation technique to the use of seeds or cuttings that germinate or root poorly. Air layers were applied root hormone to the stem branches of *Osyris lanceolata* (still attached to the plant) during Novmbere 2019, at Bazawit Hill, Nothern Ethiopia at edge of Blue-Nile River from its source at Lake Tana. Root initiation were starts after 12 weeks of the experments.The factors assessed in this experiment were the effect IBA as rooting promoter at three concentrations (0,50, 100 and 150 ppm). From the data collected it was observed 88.8% rooting were achieved from air layers in the mother plants it selef those treated by IBA hormone and the controls not responding root.Making this propagation technique is a viable alternative to the use of seed or cutting propagation. Rooting response success were influenced by application of rooting hormone of IBA, soil composition and the seasons. At a rate of 150 ppm 100% all expermented plants were intiated and primarly other than other treatmeants responded root. The significance enhancing of root making on *Osyris lanceolata* plants stem via air layers is linked to the advantage of more rooting hormone concentartion.

## 1. Introduction

*Osyris lanceolata Hochst & Steudel*. commonly known as African Sandalwood. It is a shrub or small tree growing to a height of one to seven meters (Walker, 1966; Palmer and Pitman, 1972; Mbuya et al., 1994).The species belongs to the Santalaceae family and is among the sandalwood species known for producing fragrant-scented wood from which sandal wood essentialoil is extracted. Sandalwood oil is used in various cosmetics and fragrance industry and has gained popularity in medicine (Walker, 1966; Iyenga, 1968; Srinivasan et al., 1992). The use of *Osyris lanceolata* became popular in the early 1900s following a decline in availability of Indian sandalwood, Santalum album (Eggling and Dale, 1962; Walker 1966; Iyenga, 1968; Srinivasan et al., 1992). Other species that had been considered as alternatives include S. spicatum (Errickson etal., 1973; Srinivasan et al., 1992), S. lanceolatumandS. yassi (Walker, 1966; Srinivasan et al., 1992).Since *Osyris lanceolata* was identified as a suitable alternative, it could only be found in natural stands in East Africa (Burgess et al., 1998; Mbuya et al., 1994; Ruffo et al., 2002). *Osyris lanceolata* is distributed in African countries such as Tanzania and Kenya frequently found in arid to semiarid areas, primarily on stony and rocky soils (Kokwaro, 2009) or occasionally in rocky sites and along the margins of dry forests, evergreen bushland, grassland, and thickets at an altitude range of 900-2250 m above sea level (Giathi *etal.*, 2011; Kamondo, 2012).

In East African countries, *Osyris lanceolata* constituted an important source of medicine (Mwang’ingo *et al.*, 2006). A decoction of the bark and root is considered to be useful for treating diarrhoea, gonorrhea, chronic mucus infections, and urinary diseases (Teklehaimanot, 2004; Kokwaro, 2009), a decoction of the bark in boiling water is used to treat candidiasis and related fungal infections (Masevhe *etal*., 2015) while the essential oil extracted from the bark is used to treat diarrhoea, chest problems, and joint pains. Fibers from the roots are used in basket making while the strong red dye from the bark and roots is used in skin tanning (MBUYA *etal.*, 1994). Since Osyris lanceolata is an evergreen tree with long flowering periods, it is a good forage plant (Fichtl and Adi., 1994). Several communities in Kenya also use Osyris lanceolata to produce dyes, to treat various ailments, and to brew herbal tea (Kamondo *etal*., 2012).

How ever, Osyris lanceolata is critically endangred since propagation of by seeds is difficult due to a limited supply and availability of seed at the right time (being a dioecious species, the spatial distribution of trees affects the reproductive outcome (Mwang’ingo *et al.*, 2006), storage difficulties and thus poor germination (Mbuya *etal.*, 1994). Consequently, several interventional measures are required to conserve *Osyris lanceolata*. A study by (Kokwaro, 2009) on the storage and pre-sowing treatments on seed germination demonstrated that the testa covering the embryo plays a significant role in limiting germination by restricting gas and water entry and also acts as a mechanical barrier to embryo growth. However, complete removal of the testa and soaking the zygotic embryo in hot water enhanced seed germination by 66.5%, shortened the time to seedling emergence and promoted early seedling growth (Mwang’ingo *et al.* (2006). Stem cuttings (8-10 cm long with 3-4 leaves from young trees or seedlings) could be induced to root with a maximum of 15% rooting when dipped first in a fungicide (Bavistin) for 5 min, then in 1% indole-3-butyric acid (IBA) for 6 h, but “this concentration could be increased to 32.5% when 75% of the original leaves were left intact” (Giathi *etal.*, 2011). In the same study (Giathi *etal.*, 2011), the choice of substrate was shown to affect the rooting ability of cuttings, with 30% of cuttings rooting in vermiculite, which was superior to sand, vermiculite + sand (1:1), activated coconut peat and peat. Teklehaimanot et al. (2004) used 50-150 mg/L IBA to enhance root production in young stem cuttings collected in early spring. (Mwang’ingo *et al.*, 2006) initiated air layers that were left on parent trees for eight weeks and watered every two days to allow root initiation with the help of three concentrations (50, 100 and 150 mg/L) of IBA during February, June, September, and December: 50 mg/L IBA was optimum for root initiation and June to September was best for air layering with about 80% rooting success after potting plants in sand, forest soil and animal manure (2:1:1) and fertilizing with 5 g/container of NPK (nitrogen: phosphorus, potassium) fertilizer (Machua *et al.*, (2009)) achieved 60% rooting success through air layering.

In order to restore the previous stocks of san-dalwood species in its natural stands, conventional breeding of sandalwood for introgression of new genetic in-formation can be used. However, it is an expensive and difficult task because of its long generation time, sexual incompatibility and heterozygous nature (Rugkhla and Jones.,1998). Hence, this research was attempted to addreses the endangred and economical important *Osyris lanceolata* species via the approache of air layer by injecting IBA rooting hormone at different concentration in its stem of mother trees with out detaching.

## 2. Material and Method

### 2.1. Studied area

Experments were conducted in Novmbere 2019, at Bezawit Hill, near to Blue-Nile River at Bahrdar town. It is far two-and-a-half kilometres south of from the Martyrs Memorial, Bahrdar town, Northen Ethiopia.Bezawit Hill was a palace of Haile Selassie.

### 2.2. Experimental design and treatments

An experiment for developing rooting and production of uniform plant through air layering in the stems of *Osyris Lanceoleta* plant were under take. Air layering involves rooting of stem branches without removing them from the mother plant. Experiments were carried out during dry season (Novmber,2019-March 2020)

Three factors in relation to new root formation were investigated. The effect of the season at which IBA hormone were inected in Novmber 2019 and the effect of rooting hormone concentration (lndole-3-Butyric Acid) application at concentrations of 0, 50ppm, 100ppm and 150 ppm were expermented and the soil composition. The control set (0 ppm) was treated with distilled water. For hormone injection to the stem of *Osyris lanceoleta* plant, it was applied by randomly selected four trees and remove each 10 cm length barks from its node and grild its stem as shown in **figure 1** then inject 5ml hormone for each treatments and 5ml distiled water for control by syringe from the tip of the stem bud like as shown **figure 2** since in this area there is a potent cell which is ready to develop different organs. Then, wrapped this grild area by 100 gm of sand(30%), redish soil(25%) and forest soil(45%) via poly ethline tube and seald by parafilm and alumnime foil after feaching water as above shown in **figure 3**. Each lndole-3-Butyric Acid (IBA) hormone concentrations were prepared by dissolving the proposed IBA powder with appropriate amount distilled water (50 mg IBA hormone per 1 litre for 50 ppm,100mg IBA hormone per 1 litre for 100 ppm and 150 mg IBA hormone per I Litre) to make a desirable concentration.

**Figure 1.**
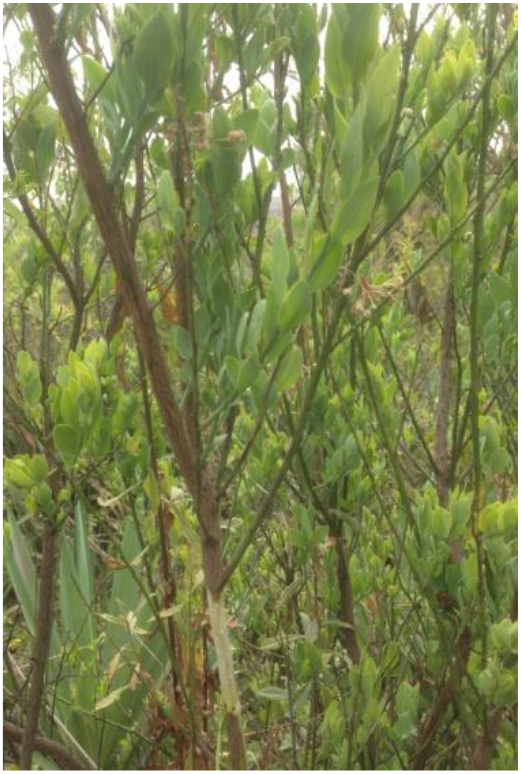
Osyris Lanceoleta in Bazawit Hill Removed barks and grild

**Figure 2.**
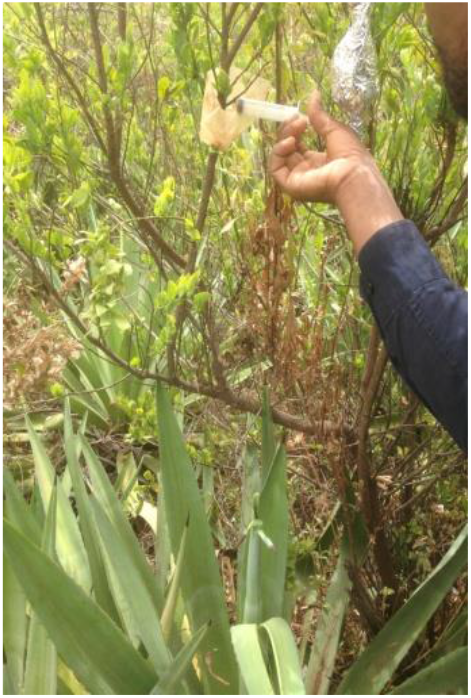
Injecting the grild area by IBA

**Figure 3.**
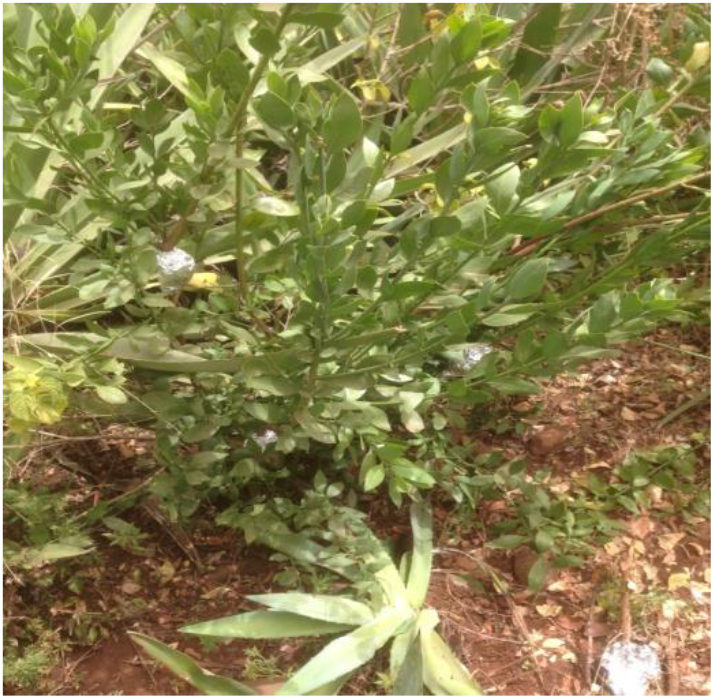
Covered the stem by aluminum foil

**Figure 4.**
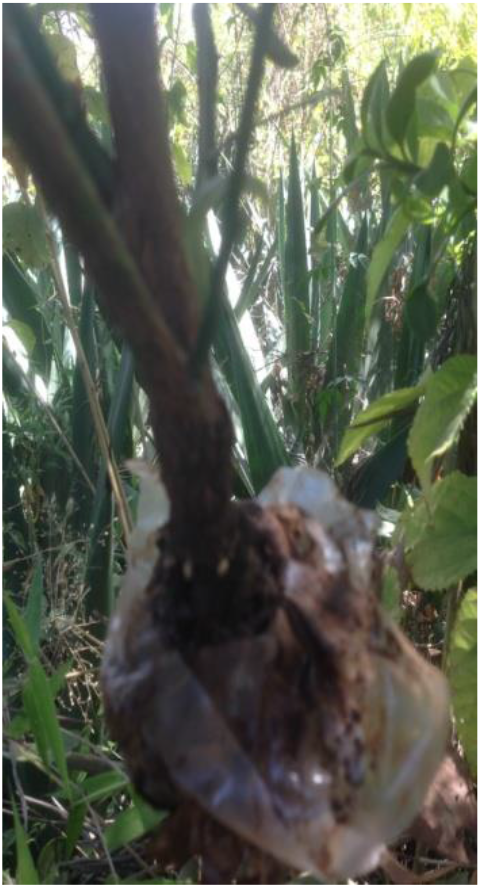
Intiated New roots by air layer propagation after 12 weeks of experment by 100 ppm concentration of IBA hormone

**Figure 5.**
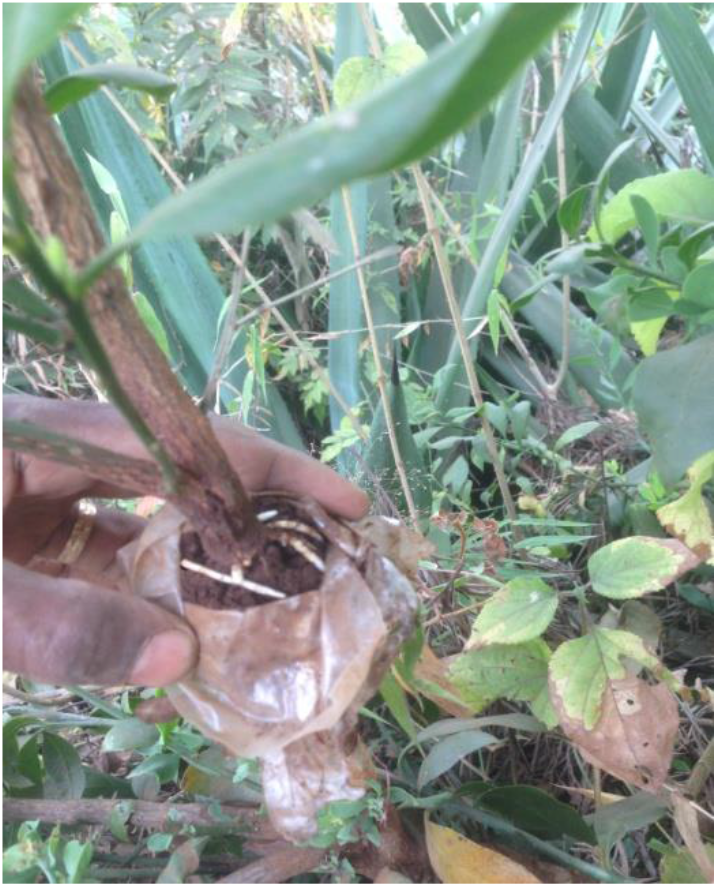
After 16 weeks of the air layring experment by 100 ppm concentration

### 2.3. Experimental management

The air layers experments were left on the parent trees for 20 weeks to allow root initiation. During this period, air layer experment were watered every four days and inspected every four week for showing weather it respond root or not. Each air layer treatment were replicated three times in each 50,100, 150 ppm IBA concentration and for control 0ppm or distiled water were applied at Bzawit hill, Bahrdar.

## 3. Results and Discusion

New root success were achieved in air layring approches that were conducted in Novmbere, 2019 at Bazawit Hill. The firist root responding time were at 12 weeks or 4 monthes of the experment and root responding time of each treatment were differed. However, 88.8 % of all the treatead plants by IBA hormone were formed root after 16 weeks of the experment.Among all treatments only at 50 ppm experment one stem plants not responding rooting after 16 weeks of the experment.

The combined effect between the experment conducted season, soil composition which wre use for wraped the grild area of the steams and IBA concentration showed a good root initiation in the present study of air layer propagation technology.The treated stems of *osyris lanceoleta* with IBA 150 ppm initially showed root respond other than other treatmeants at 12 weeks of experment. Also, after 16 weeks of the experment 150 ppm showed more root and length as shown in **Figure 6.**

**Figure 6.**
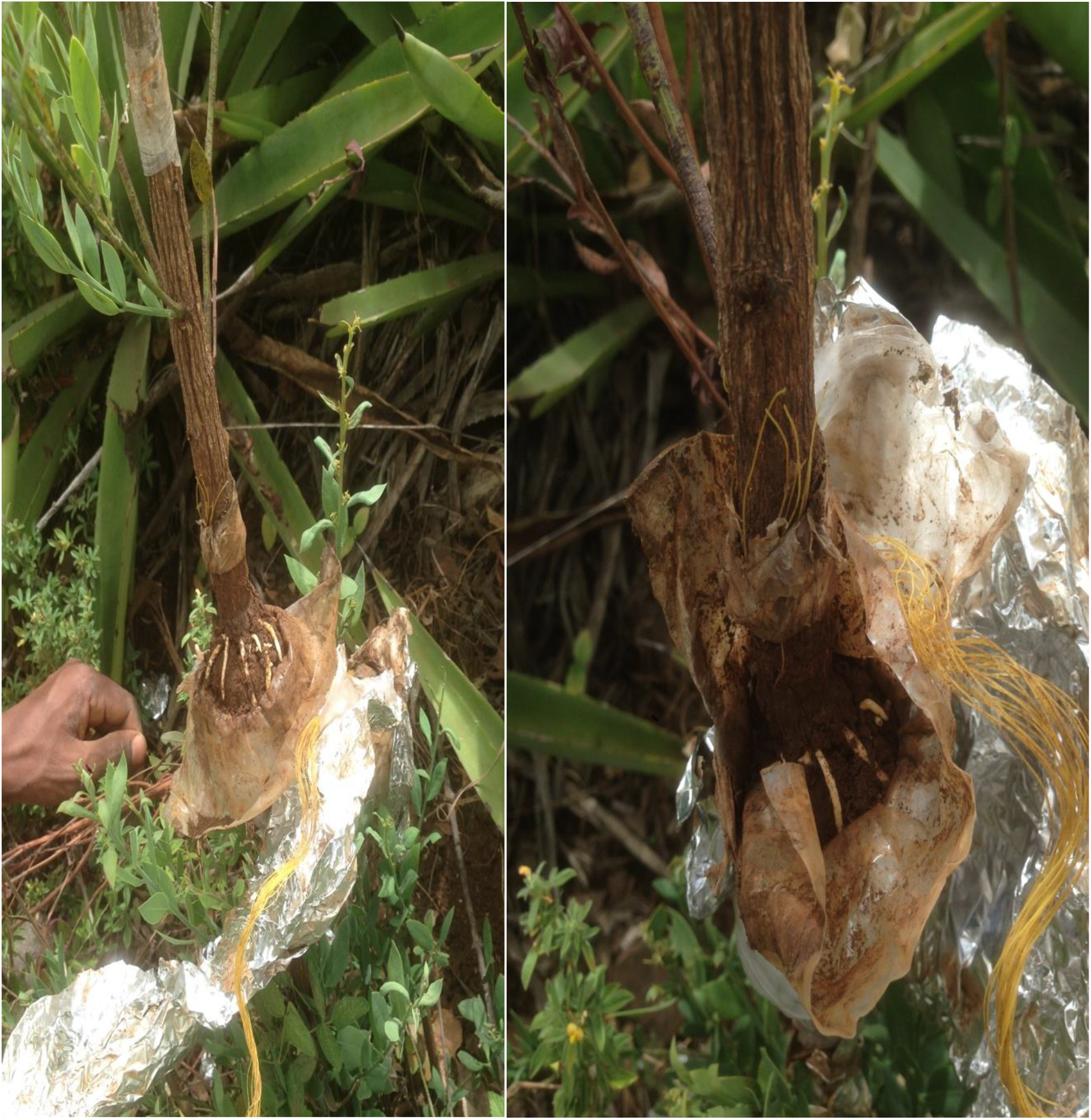
The effectes of IBA hormone at 150 ppm concentration in the stem of *Osyris Lanceoleta* after 15 weeks of the experment

The observation that auxin or IBA application initiating root and root development of the air layers is not surprising as it is commonly reported to do so in other woody plants (Howard and Harrison-Muray,1985; Leakey et al., 1982). IBA is known to influence adventitious root formation by stimulating cell division and thus increasing the ability of the airlayer to root (Hartmann and Kester, 1997; CHAUHAN et al., 1994). The variation in the amount of auxin hormone required to induce rooting for a particular season could be attributed to the differences in the amountof endogenous hormone and other associated root co-promoters, which are also known to vary with season(Nand and Anand 1970; Cheffins and Howard, 1982;Joshi et al., 1992) In the present study, 100 and 150 ppm of IBA were 100 % sucessfully root formed on *Osyris lanceoleta* Plant stem.

## 4. CONCLUSIONS AND RECOMMENDATIONS

The studied time of this experment for initiation of root on stem of *Osyris Lanceoleta* were best. Indole-3-Butyric Acid (IBA) were also enhancing the rooting ability and development of air layers with the studied time of the experment. For actively root formation IBA concentration up to 150 pp are required.

Air layering produced better success compared with other propagation studies since *Osyris lanceolata* is naturally have low regeneration and very limited seeds quality has been reported in diffrent scientfic research studies, hence, multiplication of the species through air layering is an alternative and effective option for producing clonal plants.Further, well established plants could be obtained within a short time. The method is also inexpensive and easy to perform, and therefore, will be acceptable to the locals nursery and those involved in the forestry sector.

Finally, any researchar which have intersted to do more research on *Osyris lanceolata*, it can get this plant in Ethiopia from arid to semiarid areas, on stony and rocky soils and along the margins of dry forests, evergreen bushland and grassland. Among the forest land which is *osyris lanceoleta* scateredelly found in Bazawit Hill, Abay Berha /Deserte, Debere Libanose Monistery, Taragedame, Alem Saga forest, Neche Sare National Parak and mostelly in Omo valley.

## 5. Acknowledgments

I am highly acknowlede Ethiopian Enviroment, Forest and Climate Change Commisions by finacing this research via UNDP Institutional Strengthening for the Forest Sector Development Program.

